# High-throughput Production of Diverse Xenobiotic Metabolites with P450-transduced Huh7 Hepatoma Cell Lines

**DOI:** 10.1101/2022.03.12.484071

**Authors:** Choon-Myung Lee, Ken H. Liu, Grant Singer, Gary W. Miller, Shuzhao Li, Dean P. Jones, Edward T. Morgan

**Affiliations:** Department of Pharmacology and Chemical Biology, Emory University School of Medicine, Atlanta, Georgia 30322; Clinical Biomarkers Laboratory, Department of Medicine, Emory University, Atlanta, Georgia 30322; Department of Environmental Health Sciences, Columbia University Mailman School of Public Health, New York, New York 10032; The Jackson Laboratory for Genomic Medicine, Farmington, Connecticut 06032

## Abstract

Precision medicine requires methods to assess drug metabolism and distribution, including the identification of known and undocumented drug and chemical exposures as well as their metabolites. Recent work demonstrated high-throughput generation of xenobiotic metabolites with human liver S-9 fractions and detection in human plasma and urine. Here, we developed a panel of lentivirally transduced human hepatoma cell lines (Huh7) that stably express individual cytochrome P450 (P450) enzymes and generate P450-specific xenobiotic metabolites. We verified protein expression by immunoblotting and demonstrated that the cell lines generate P450-specific metabolites from probe substrates. To increase analytical throughput, we used a pooling strategy where 36 chemicals were grouped into 12 unique mixtures, each mixture containing 6 randomly selected compounds, and each compound being present in two separate mixtures. Each mixture of compounds was incubated with 8 different P450 cell lines with cell extracts analyzed at 0 and 2 h. Extracts were analyzed using liquid chromatography-high resolution mass spectrometry. Cell lines selectively metabolized test substrates, with pazopanib metabolized by CYP3A4 and CYP2C8 cells, bupropion by CYP2B6, and β-naphthoflavone by CYP1A2 for example, showing substrate-enzyme specificity. Predicted metabolites from the remaining 33 compounds as well as many unidentified *m/z* features were detected. We also show that a specific metabolite generated by CYP2B6 cells, but not detected in the S9 system, was identified in human samples. Our data show that incubating these cell lines with chemical mixtures accelerated characterization of xenobiotic chemical space, while simultaneously allowing for the contributions of specific P450 enzymes to be identified.

**Significance statement:** High resolution mass spectrometry enables the identification of exposures to drugs and other xenobiotics in human samples. This paper demonstrates a workflow for high throughput production of xenobiotic metabolites using a panel of engineered cytochrome P450-expressing hepatoma cells. Active substrate-enzyme pairs can be identified using this workflow and generated metabolites can be used as surrogate standards to validate xenobiotic detection in humans.

## Introduction

Widespread use of high resolution mass spectrometry (HRMS)-based metabolomics methods shows that many unidentified mass spectral features are significantly associated with human diseases, with many features likely arising from undocumented drug, dietary supplement and environmental chemical exposures (Jones, 2016; Uppal et al., 2016). Despite advancements in instrumentation and bioinformatic tools for chemical annotation (Uppal et al., 2017; Blazenovic et al., 2019) and prediction of biotransformation (Djoumbou-Feunang et al., 2019), the lack of authentic standards for xenobiotic metabolites is an obstacle for identification of xenobiotic exposures. Developing a high-throughput platform for generation of biotransformation products for thousands of xenobiotic chemicals would establish experimentally validated sets of xenobiotic metabolites that could be used for identification of undocumented exposures in people.

We recently used human liver S9 fractions to develop an *in vitro* high-throughput biotransformation platform coupled with HRMS, showing that generated xenobiotic metabolites can be used as reference mixtures to identify xenobiotic exposures in human blood and urine (Liu et al., 2021). However, one disadvantage of using S9 fractions for biotransformation is that not all xenobiotic metabolizing enzymes are equally expressed. As a result, the metabolic contributions of enzymes that are less highly expressed or that have lower specific activity may be missed. In the present study, we used lentiviral transduction to develop a panel of cytochrome P450 (P450, CYP)-expressing Huh7 hepatoma cell lines, each expressing a specific P450. P450s are a gene superfamily of monooxygenases that are universal in all living systems. Mammalian P450s are involved in oxidation of many substrates including fatty acids, steroids, and xenobiotics. Although there are 18 families with 57 CYP genes in humans, CYP families 1 through 3 are mainly involved in xenobiotic metabolism such as drugs, food nutrients, environmental chemicals, etc. (Nebert et al., 2013). Moreover CYP1, 2, 3 families are involved in metabolism of more than half of clinically available drugs (Cerny, 2016). We developed these cell lines as a complement to the generalized S9 system to allow for simultaneous production of xenobiotic metabolites and identification of specific enzyme(s) that are capable of producing them. The identification of substrate-enzyme pairs would allow specific P450 metabolites of interest to be scaled-up for additional characterization and facilitate the identification of xenobiotic exposures in human samples.

In validation and testing, we recognized LC-HRMS throughput as a bottleneck to large-scale characterization of xenobiotic metabolites, and so we tested a mixtures metabolism workflow with these P450 cell lines to address this limitation. 36 chemicals were grouped into 12 mixtures of 6 randomly selected compounds, with each compound present in two separate mixtures. These 12 individual mixtures were incubated with 8 different cytochrome P450-expressing Huh7 cell lines, with cell extracts collected and analyzed using LC-HRMS. Our data show that adoption of this workflow increased analytic throughput by a factor of six, and also demonstrate the formation of unexpected metabolites in a cell-line specific manner. These results could be used to identify particular xenobiotic exposures in human samples.

## Materials and Methods

### Materials

Acenaphthene, acetaminophen, auranofin, β-naphthoflavone (β-NF), benzo[a]pyrene, bupropion, caffeine, chlorzoxazone (CLZ), nicotine, tolbutamide, trifluralin, naphthalene, coumarin, fluoranthene, 1,2-diphenylhydrazine, ethylbenzene, 2,4,6-trichlorophenol, estrone, 1,1,2,2-tetrachloroethane, N-nitrosodipropylamine, 2-methylnaphthalene, endrin aldehyde, azinphos-methyl, dexamethasone, Geneticin (G418), pazopanib, S,S,S tributyl-phosphorotrithioate, bromodichloromethane, ethion, benzo[b]fluoranthene, bis(2-ethylhexyl)phthalate, crizotinib, 1,2-dichlorobenzene, o,p’-dichlorodiphenyldichloroethane (DDD), omeprazole, pazopanib, polybrene, warfarin and HPLC grade acetonitrile were purchased from Sigma-Aldrich (St. Louis, MO). S,S-hydroxy and R,R-hydroxy bupropion were purchased from Cayman Chemical, Ann Arbor, MI) and vandetanib was from AstaTech (Bristol, PA). Blasticidin was obtained from InvivoGen (San Diego, CA). Rabbit anti-V5 (V8137), mouse anti-glyceraldehyde 6-phosphate dehydrogenase (MAB374), and mouse anti-actin (A2228) antibodies were purchased from Sigma.

### Generation of P450 expressing cell lines and Immunoblotting

Human hepatoma Huh7 cell lines individually expressing CYP1A2, 2A6, 2B6, 2C8, 2C19, 2D6, 2E1, or 3A4 with a C-terminal V5 tag were generated as described in previous studies (Cerrone et al., 2020)}(Park et al., 2017; Lee et al., 2020). The lentiviral vector pLX304-2D6 (cat# EX-A3525-LX304) was purchased from GeneCopoeia (Rockville, MD). The other lentiviral vectors (pLX304-CYP) were obtained from the DNASU plasmid repository center. pLX304 plasmids with cloned V5-tagged P450s were transfected together with a second-generation lentiviral packaging system consisting of pMD2.G and psPAX2 plasmids into HEK293T cells to generate virus particles. 48 and 72 h after transfection, media containing virus particles were collected and filtered through a 0.45 µm filter, and stored at −80°C. Huh7 cells were infected with viral media containing 10% fetal bovine serum (FBS)/1% penicillin/streptomycin (PS)/Dulbecco’s Modified Eagle Medium (DMEM) and polybrene (8 μg/ml) to enhance transduction. Cells were selected with 10 μg/ml blasticidin beginning at 24 h after infection.

To confirm the expression of the individual P450s, the cells were grown to 95%–100% confluence and the proteins were extracted with cell lysis buffer. The total cell lysates were collected and centrifuged at 12,000 x *g* for 5 min and then the supernatant was collected and prepared for SDS-PAGE. Immunoblotting was carried out using anti-V5 antibodies (1:5000; Sigma) as described previously (Cerrone et al., 2020).

### P450 activity time course assays

The eight P450 cell lines were plated on 96-well plates in 10% FBS/1% PS /10 μg/mL blasticidin/DMEM media until they reached 100% confluency. After media were removed, 50 μL of media containing a mixture of seven P450 substrates (20 μM each of 7-ethoxyresorufin (CYP1A2), coumarin (CYP2A6), bupropion (CYP2B6), amodiaquine (CYP2C8), omeprazole (CYP2C19 and 3A4), dextromethorphan (CYP2D6), and chlorzoxazone (CYP2E1) were added to the wells and incubated for the indicated time. The incubations were stopped by addition of 150 μL of acetonitrile, mixed by repetitive pipetting, and centrifuged at 3000 x g for 5 min. The supernatants were transferred into 96-well liquid chromatography (LC)-ready plates for LC-HRMS analysis.

### Pooling strategy to improve throughput for xenobiotic metabolite generation

36 chemicals were grouped into 12 separate mixtures at a final concentration of 20 μM in 1% dimethylsulfoxide. Each pool contained 6 randomly assigned compounds (so that each chemical would be present in 2 distinct mixtures) in P450 assay buffer (1 mM Na2HPO4, 137 mM NaCl, 5 mM KCl, 0.5 mM MgCl2, 2 mM CaCl2, 10 mM glucose, and 10 mM Hepes, pH 7.4 (Donato et al., 2004)). The eight cell lines expressing individual P450 enzymes were cultured in 10% FBS/1% PS/10 /10 μg/mL blasticidin/DMEM media until they reached 100% confluency on 96-well plates. After removing media, each cell line was incubated with 50 μL of each of the 12 substrate pools in P450 assay buffer in two 96-well plates for 0 and 2 h resulting in 192 samples for analysis. Reactions were terminated by addition of 150 μL of acetonitrile and processed as described above for LC-HRMS.

### LC-HRMS

10 μL aliquots of sample extracts were analyzed using liquid chromatography coupled to Orbitrap-based MS analysis (Dionex Ultimate 3000, Thermo Scientific Fusion or Thermo Scientific High-Field Q-Exactive). The chromatography system was operated in a dual pump configuration which allows sample analysis on one column with a second column undergoing flushing and reequilibration in parallel. Sample extracts were injected and analyzed using hydrophilic interaction liquid chromatography (HILIC) with electrospray ionization (ESI) operated in positive mode and reverse phase (C18) chromatography with ESI operated in negative mode. Analyte separation for HILIC was accomplished using a Waters XBridge BEH Amide XP HILIC column (2.1 mm x 50 mm, 2.5 μM particle size) and eluent gradient (A = water, B = Acetonitrile, C = 2% Formic acid) consisting of an initial 1.5 min period of 22.5% A, 75% B, 2.5% C, followed by a linear increase to 75% A, 22.5% B, 2.5% C at 4 min and a final hold of 1 min. C18 chromatography was performed using an endcapped C18 column (Higgins Targa C18 2.1 mm x 50 mm, 3 μM particle size) and gradient (A = water, B = acetonitrile, C = 10 mM ammonium acetate) consisting of an initial 1 min period of 60% A, 35% B, 5%C, followed by a linear increase to 0% A, 95% B, 5% C at 3 min and held for the remaining 2 min. For both methods, mobile phase flow rate was held at 0.35 mL/min for the first 1 min, increased to 0.4 mL/min for the final 4 min. Mass spectrometers were operated at 120,000 resolution and data were collected from 85-1,275 *m/z* for MS1 analysis with data-dependent MS/MS analyses performed at 60,000 resolution for MS1 and 30,000 resolution for MS2 using an inclusion list for predicted xenobiotic metabolites.

### Identification of m/z features

Raw data were processed in mzMine2 (Pluskal et al., 2010). Extracted data were organized into a feature table data file where each ion feature is represented by its characteristic *m/z*, retention time, and associated peak intensity. Lists of accurate mass *m/z* ions for expected metabolites (M+H for HILIC+ and M-H for C18-) were generated in three complementary ways: from the literature, using Biotransformer (Djoumbou-Feunang et al., 2019) and by subtracting the exact mass of the parent compound from observed masses that increased with time and looking for discrete masses that corresponded to known biotransformations (demethylation, hydroxylation, etc.). These target lists were used to search the feature table for biotransformation products for each precursor compound and metabolite peak intensities increased at 2 h relative to 0 h time points.

## Results

### Expression of active, tagged P450s and characterization of their substrate specificities

After selecting lentivirus transformed cells with blasticidin, we carried out immunoblotting with an anti-V5 antibody to confirm the expression of P450 proteins (Fig. 1). All CYP cell lines expressed the corresponding proteins with the C-terminal V5 tag. To examine the functionality and substrate specificity of the expressed enzymes, all cell lines were incubated for 0, 2, 4 and 6 h with a mixture of 7 enzyme-selective substrates, and contents (cells and media) of each well were extracted and analyzed on the high-resolution mass spectrometer (Fig. 2). Huh7 cell lines, each labeled according to the specific CYP used for transfection (1A2, 2A6, 2B6, 2C8, 2C19, 2D6, 3A4) demonstrated robust substrate-selective activities (Fig. 2). The well-documented CYP1A substrate 7-ethoxyresorufin (Ghosal et al., 2003) was O-deethylated by the Huh7-CYP1A2 cell line to form resorufin (Fig. 2A) and the CYP2A6-specific substrate coumarin (Pelkonen et al., 2000) was oxidized only by the Huh7-CYP2A6 cell line to form 7-hydroxycoumarin (Fig. 2B). Huh7-CYP2B6 cells oxidized the CYP2B6-specific substrate bupropion (Hsyu et al., 1997) to hydroxybupropion (Fig. 2C). Additionally, we estimated the concentration of generated hydroxybupropion with the R,R-hydroxy bupropion standard (Fig. 3). Hydroxybupropion was generated rapidly within 4 h and continued to be produced until 24 h (Fig. 3B). At 24 hours, half of the added 10 mM buproprion had been converted to hydroxybupropion (Fig. 3B), showing the utility of this cell line to generate xenobiotic metabolites in large quantities. The CYP2B6 cell line also produced dihydroxybupropion from bupropion, although it was observed at 10-fold lower relative abundance compared to hydroxybupropion (Fig. 3C). Dihydroxybupropion was previously detected in human urine, but its metabolic source was unknown (Petsalo et al., 2007). Furthermore, we observed dihydroxybupropion in mice following oral gavage of bupropion (Fig. 4A), and in a plasma sample from a patient with documented bupropion use in which we had previously identified hydroxybupropion and hydrobupropion (Liu et al., 2021). The retention times and ion fragmentation spectra for dihydroxybupropion detected in the patient matched that of the CYP2B6 generated dihydroxybupropion (Fig. 4B).

**Fig 1.**
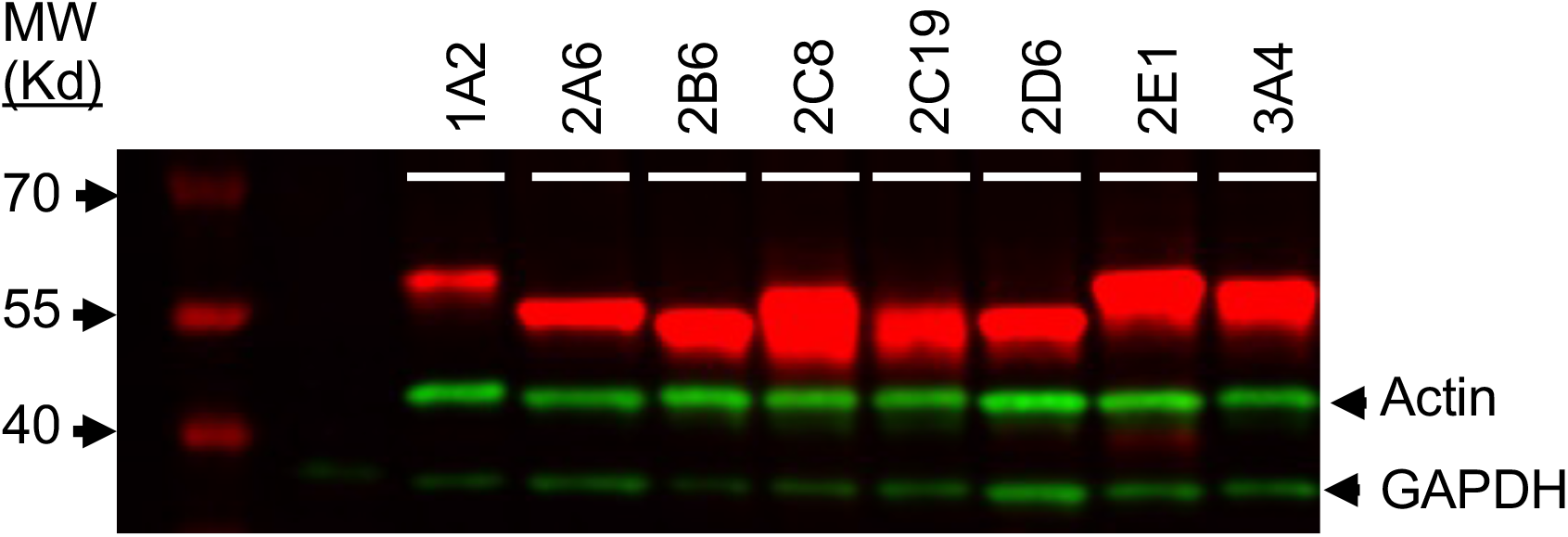
Immunoblotting of P450 expressing cell lines. Equal volumes of total cell extracts from the indicated cell lines were separated on SDS-PAGE and blotted with anti-V5, anti-actin, and anti-GAPDH primary antibodies.

**Fig 2.**
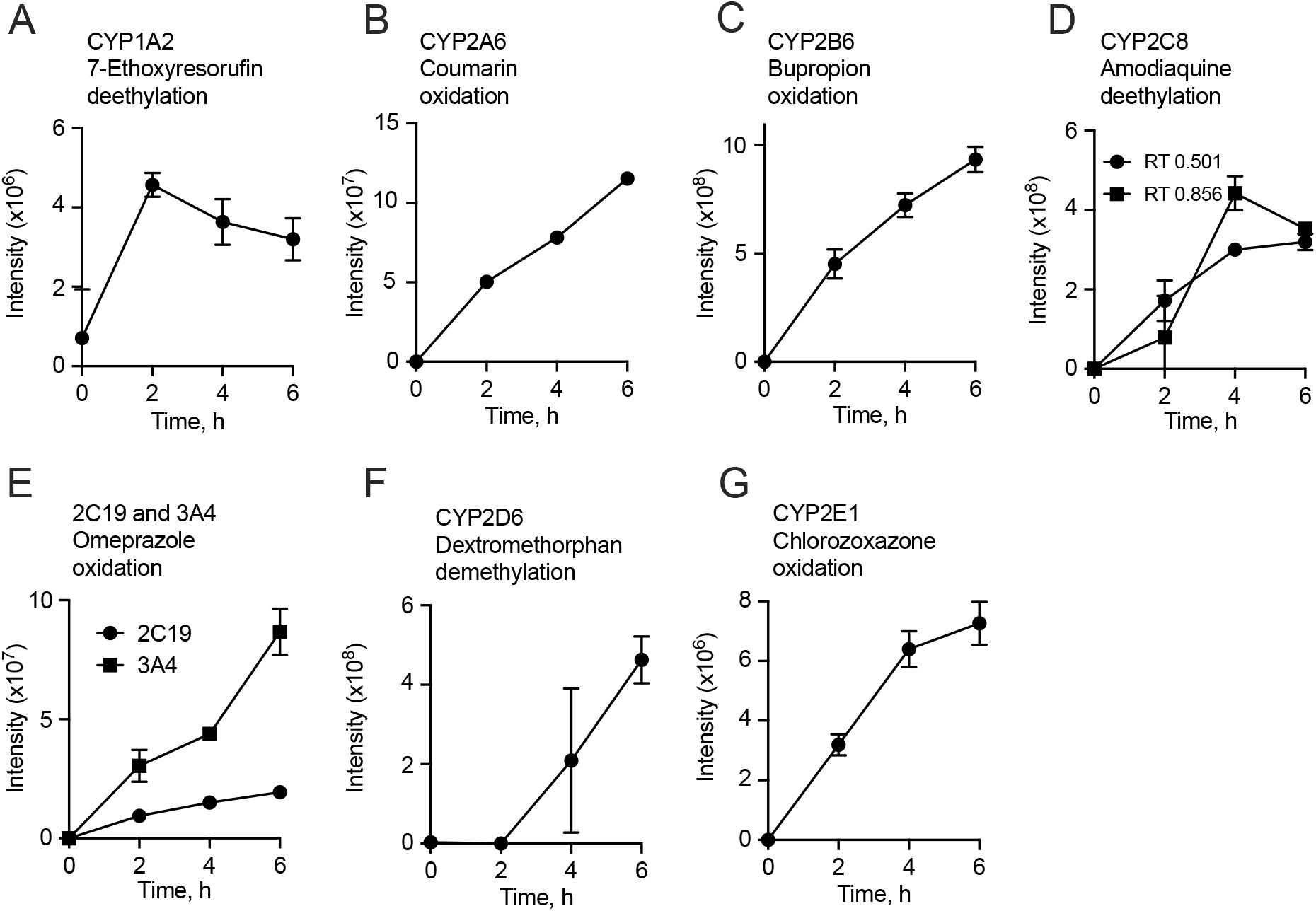
Cell line and substrate specificity. A mixture of 7 substrates at 20 μM final concentration were incubated with each P450-expressing cell line for the indicated times and then quenched with 3 volumes of acetonitrile (ACN) and then analyzed by LC-HRMS (Q Exactive HF mass spectrometer) Only detected signals were plotted. The predicted product masses was selected by 3 criteria: 1) mass (*m/z*) tolerance should be within 0.0005, 2) masses should have the same retention time, and 3) the signal should increase over time. All signals were detected in HILIC positive mode except chlorzoxazone in *panel G* where the signal was detected in C18 negative mode. A, ethoxyresorufin deethylation (BROD) by CYP1A2 cell line; B, coumarin hydroxylation by CYP2A6 cell line; C, bupropion hydroxylation by CYP2B6 cell line; D, amodiaquine deethylation by CYP2C8; E, omeprazole hydroxylation by CYP2C19 and 3A4 cell lines; F, dextromethorphan demethylation by CYP2D6; G, chlorzoxazone (CLZ) hydroxylation by CYP2E1 (C18). CYP2E1 cell lines were incubated with only chlorzoxazone for the indicated time and analyzed on the same LC-HRMS since we were not able to detect the OH-CLZ from the mixed substrate samples. Values are means ± SD of three individual wells from the same cell preparation.

**Fig. 3.**
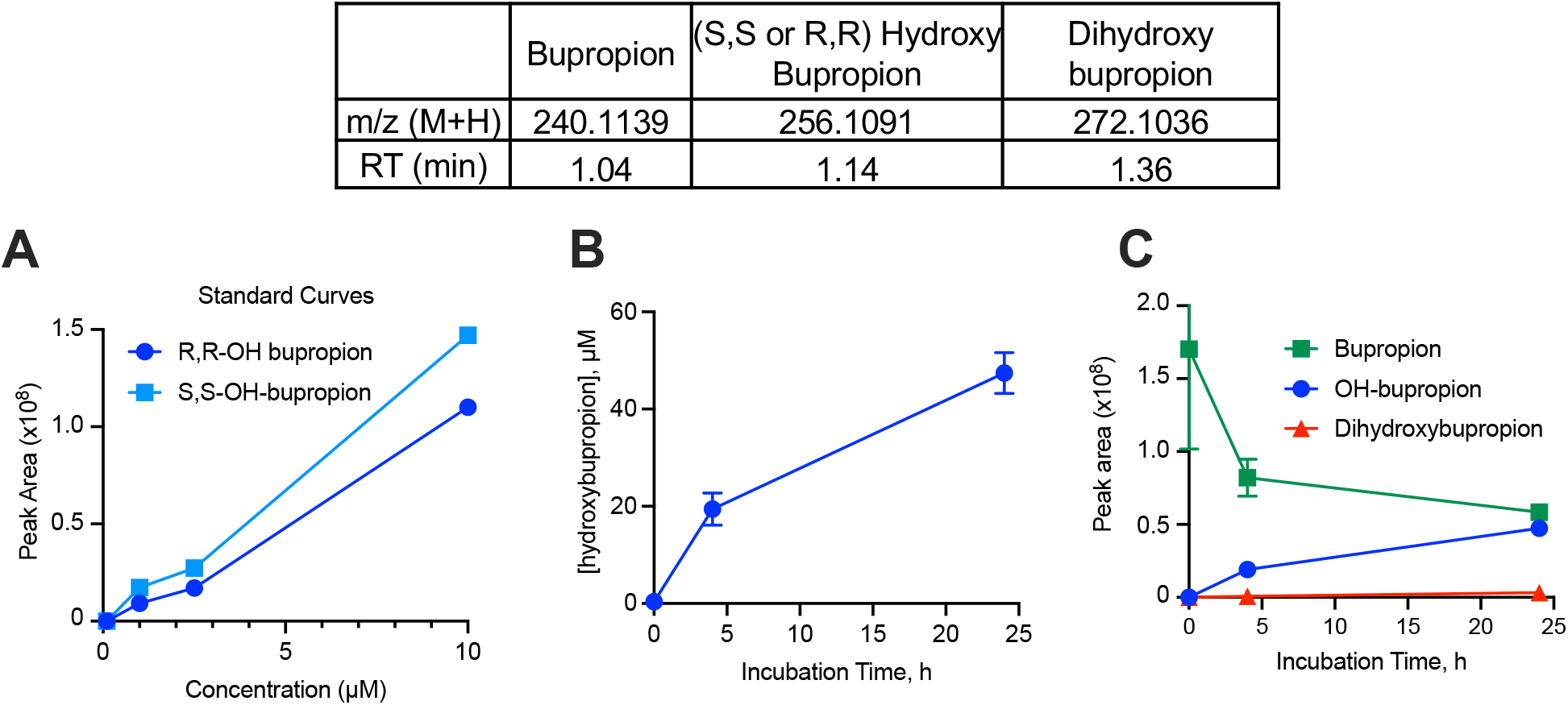
Estimation of hydroxybupropion generation by the Huh7-2B6V5 cell line. Huh7-2B6V5 cells were plated on 24-well plates at 80% confluency and then two-days later, media (10% FBS/PS/DMEM) containing 100 mM of bupropion was added for various incubation times. After incubation of substrate, 3 volumes of ACN were added and the plate was frozen and thawed. Cells were broken by pipetting and after spinning down the cell-media mixtures, supernatants were collected to vials for the HRMS analysis (Orbitrap ID-X Tribrid MS). A. Standard curve of S,S-hydroxy and R,R-hydroxy bupropion. B. Quantitation of hydroxy bupropion generation using the R,R-hydroxy bupropion standard. Values are the means ± SD of three individual wells from the same cell preparation. C. Detection of bupropion, hydroxy and di-hydroxy bupropion signals at indicated times from the same experiment as in panel B.

**Fig. 4.**
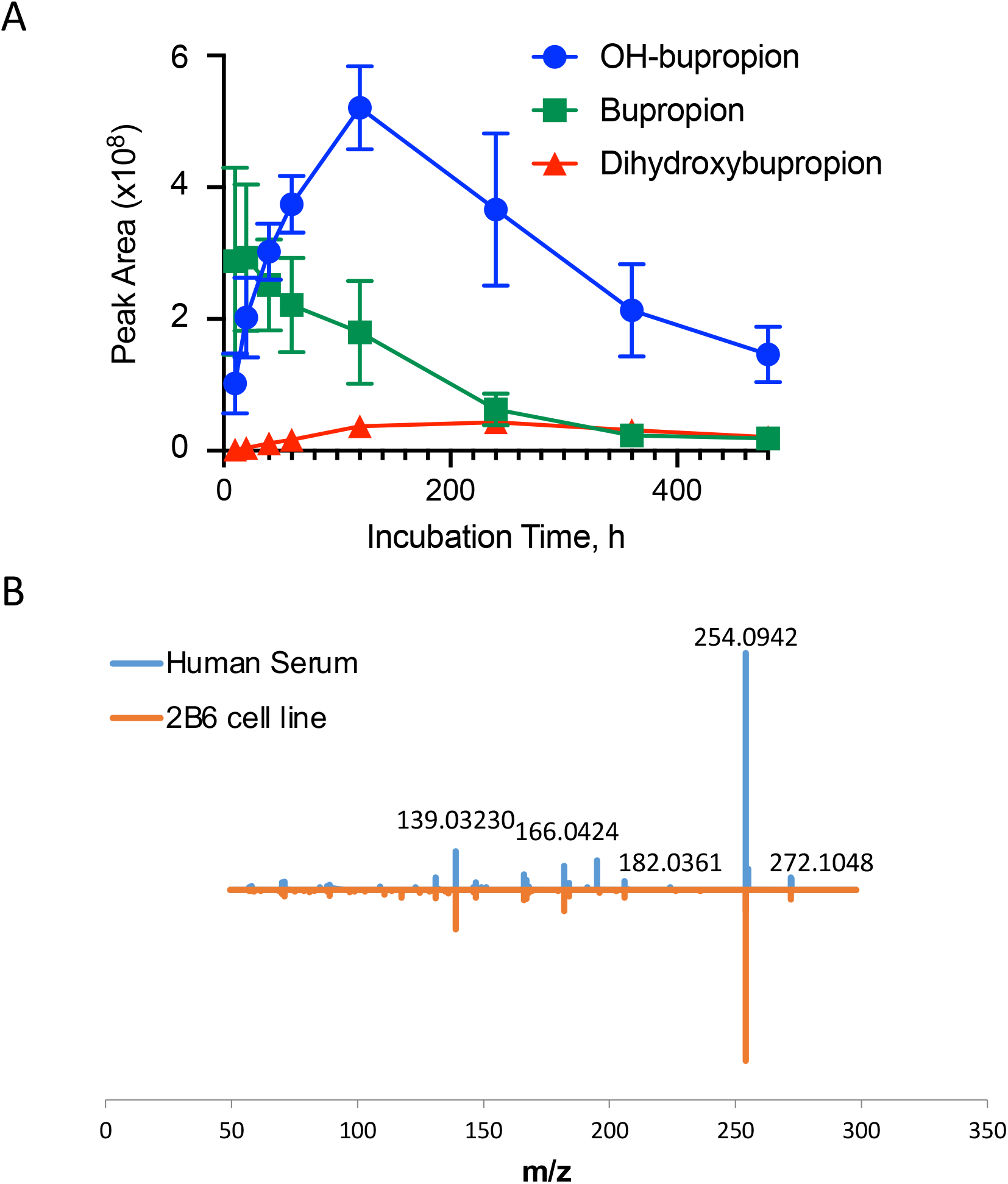
Pharmacokinetics of bupropion and its metabolites, hydroxy and dihydroxybupropion from C57BL mouse after bupropion oral gavage. A. The data for bupropion was published previously (Mimche et al., 2019), and the feature tables were re-analyzed to identify hydroxy and dihydroxybupropion based on their retention times, m/z values and MS^2^ spectra. Values are means ± SD of 5 mice in each group. B. Mass fragmentation spectra of 272.1048 m/z (dihydroxybupropion) in 2B6 cell line (red) and in human sample (blue) with documented bupropion use

A well-established CYP2C8 substrate amodiaquine (Li et al., 2002) was N-deethylated by the CYP2C8 cell line producing two forms of desethylamodiaquine (RT, 0.501 and 0.856 min) in a time-dependent manner (Fig. 2D). Omeprazole was oxidized by two cell lines, CYP2C19 and CYP3A4 as expected (Yamazaki et al., 1997; Li et al., 2005) (Fig. 2E). However, CYP3A4 cells were 4.5 fold more active than CYP2C19 cells. Dextromethorphan is well a known probe drug for human CYP2D6 activity (Yu and Haining, 2001), and the CYP2D6 cell lines actively metabolized dextromethorphan to desmethyldextromethorphan (probably O-demethylation to dextrorphan) at 4 and 6 h, while no product was detected at 2h (Fig. 2F). An additional experiment showed that the CYP2D6 cell line actively demethylated dextromethorphan within 2h (Supplemental Fig. 1A), as well as generating hydroxydesmethyldextrorphan (Supplemental Fig. 1B). Our data suggest that the CYP2D6 cell line is more active in hydroxylation of desmethyl dextromethorphan than dextromethorphan because we did not detect the hydroxydextromethorphan. Our initial experiment did not detect the CYP2E1 probe drug chlorzoxazone (CLZ) parent compound or hydroxychlorzoxazone formation by CYP2E1 cell lines when using the substrate mixture. However, a second experiment using CLZ as the lone substrate showed the time-dependent generation of OH-CLZ by CYP2E1 cells detected in C18 negative mode (Fig. 2G), suggesting inhibition of activity by other drugs or drug metabolites in the mixture. The data show that these P450-expressing cell lines have robust cell line-specific (P450 specific) reactions and can be used for drug metabolism studies with some potential for interference when assayed as mixtures.

### Cell-line specific metabolite production from pooled mixtures

The above experiments using pooled probe substrates suggest that a substrate pooling strategy may facilitate efficient screening of large numbers of chemicals for metabolite production. 36 compounds were pooled randomly into 12 pools with 6 individual compounds in a pool (Fig. 5; details provided in Supplemental Fig. 2) and each pool was incubated separately with each of the 8 P450-expressing cell lines for 0 and 2 hprior to analysis by LC-HRMS. The resulting complex data (Fig. 6, Table 1) were simplified by targeted searching for predicted metabolites as well as application of data tools to search for other non-predicted products; selected examples are provided to illustrate the utility of the cell lines for generation of products to aid in metabolite identification in human metabolomics studies and also to discover an uncharacterized metabolite.

**Table 1.** Xenobiotic biotransformation products from CYP cell lines. Metabolites detected in HILIC-positive mode are shown. Targeted metabolites are named. Numbers in parentheses represent unidentified biotransformation products which were strongly associated with parent compounds and identified by the criteria stated in the text. (Pearson coefficient >0.85).

| CYP cell line | 1A2 | 2A6 | 2B6 | 2C8 | 2C19 | 2E1 | 2D6 | 3A4 |
| --- | --- | --- | --- | --- | --- | --- | --- | --- |
| Acenaphthene |  |  |  |  |  |  |  | (4) |
| Fluoranthene | (7) | (56) | (16) | (58) | (7) | (37) | (7) | (14) |
| 1,1,2,2-Tetrachloroethane | (2) |  | (2) | (1) |  |  |  |  |
| Azinphos methyl |  | (1) |  | (4) | (6) | (1) | (3) |  |
| s,s,s, tributylphosphorotriothioate | (3) |  | (2) | (1) | (2) | (3) | (4) | (2) |
| β-naphthoflavone | di-OH (3) |  |  | (1) |  |  |  |  |
| Chlorzoxazone | (2) | (1) | (1) | (1) |  |  |  | (2) |
| 1,2-diphenyl hydrazine | (1) | (1) | (1) | (3) |  | (1) |  |  |
| N-Nitrosodipropylamine | (1) |  | (1) | (2) | (3) |  |  |  |
| Dexamethasone |  | R (1) |  | R (1) |  | R (2) | R (1) | R (1) |
| Bromodichloromethane | (1) | (1) |  |  | (1) |  | (1) |  |
| 1,2-dichlorobenzene |  | (1) |  |  |  |  |  |  |
| APAP | (6) | (4) | (4) | (8) | (3) |  | (5) | (2) |
| Ethylbenzene |  |  |  |  |  |  |  |  |
| Benzo[a]pyrene |  | (1) |  |  | (1) |  |  |  |
| Bupropion |  | DA (1) | OH (1) | (2) |  | (1) | (1) |  |
| Ethion | (2) | (1) | (5) | (4) | (15) |  |  | (5) |
| Vandetanib | MK (5) | (1) | (11) | (13) | (6) | (1) | (4) | (2) |
| Naphthalene |  |  |  | (1) |  |  |  | (4) |
| Tolbutamide | (6) | (2) | (7) | (4) | OH (10) | (1) | (9) | (8) |
| 2-Methylnaphthalene | (1) |  |  |  | (1) |  | (1) | (1) |
| G418 | (1) |  |  |  | (3) | (1) |  |  |
| Benzo-b-fluoranthene | (1) |  | (1) | (2) |  |  |  |  |
| o,p,-DDD |  |  | (3) |  | (1) |  |  | (1) |
| Coumarin | (4) | (13) |  | (1) | (2) | (1) | (1) |  |
| 2,4,6-Trichlorophenol |  |  |  |  |  |  |  |  |
| Endrin aldehyde |  | (1) |  |  |  |  |  |  |
| Caffeine | (3) |  | (1) | (1) | (3) |  | (5) | (1) |
| Bis-2-ethylhexylphthalate | (3) | (6) | (5) | (2) | (3) | (3) |  | (5) |
| Nicotine | (3) | (4) | +Ch3 ?,<br>OH, di-OH,<br>MK, R? (1) |  | +Ch3 ? (1) | +Ch3 ? (1) |  | +Ch3 ? (2) |
| Trifluralin | (1) |  | (7) | (1) |  | OH, di-OH,<br>R? (6) | (1) | (13) |
| Estrone | (3) |  |  |  |  |  |  |  |
| Auranofin | (16) | (18) | (16) | (24) | (27) | (21) | (27) | (32) |
| Pazopanib | (1) |  |  | OH (2 RTs)<br>(5) |  | (2) | (2) | H, DA (4) |
| Crizotinib | (3) | 16 | (6) | (2) | (1) | (2) |  | (7) |
| Warfarin | (2) | (2) | (1) | (3) | (3) | (5) | (3) | (1) |

+CH<sub>3</sub>, methylation; DA, dealkylation; di-OH, dihydroxylation; MK, methylene to ketone; OH, hydroxylation; R, reduction.

**Fig 5.**
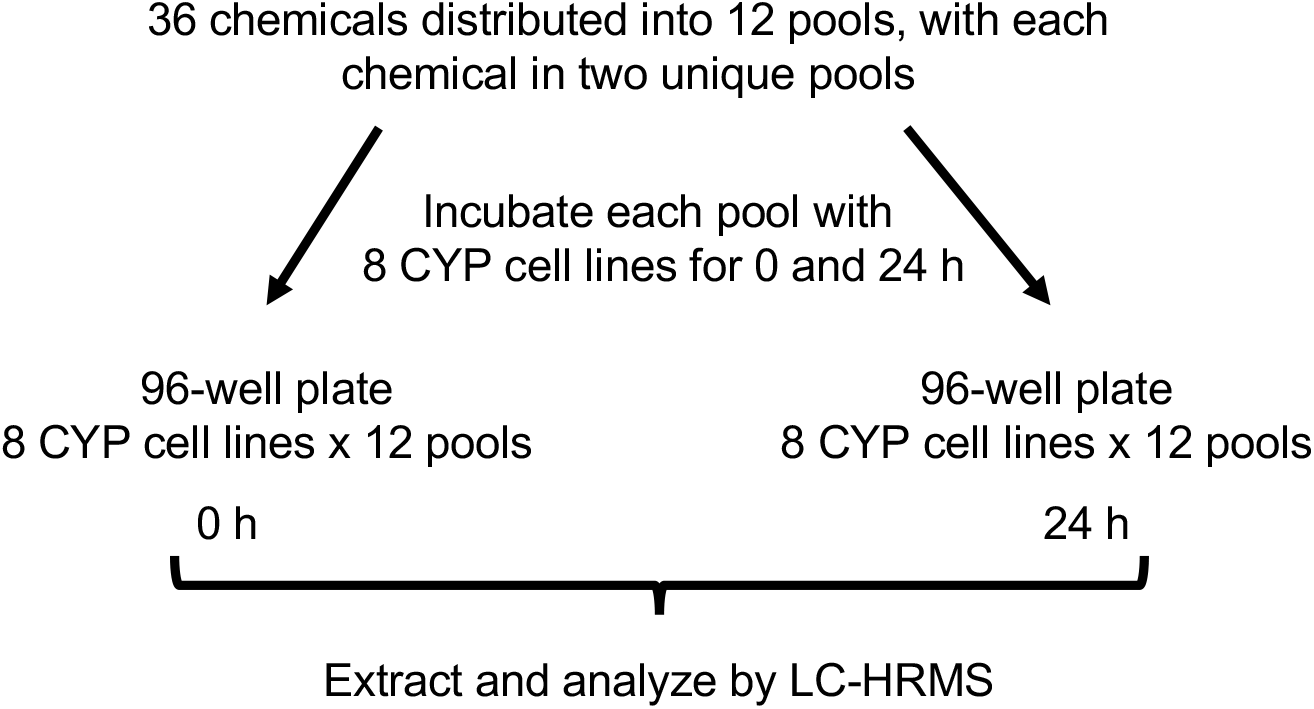
Pooling strategy. A. Pooling samples. A total of 36 compounds were grouped into 12 pools, with each pool containing a unique combination of 6 compounds and each compound being present in two separate pools. Each chemical was present at a final concentration of 20 μM in 0.1% dimethylsulfoxide. B. Experimental design. After each pool was incubated with 8 P450-expressing cell lines for 0 and 2 h on two 96-well plates, respectively, reactions were quenched with 3 volumes of ACN and then analyzed by LC-HRMS (Orbitrap Fusion(tm) Tribrid(tm) Mass Spectrometer). For additional details concerning cell lines and xenobiotic pools, see Supplemental Fig. 3.

**Fig 6.**
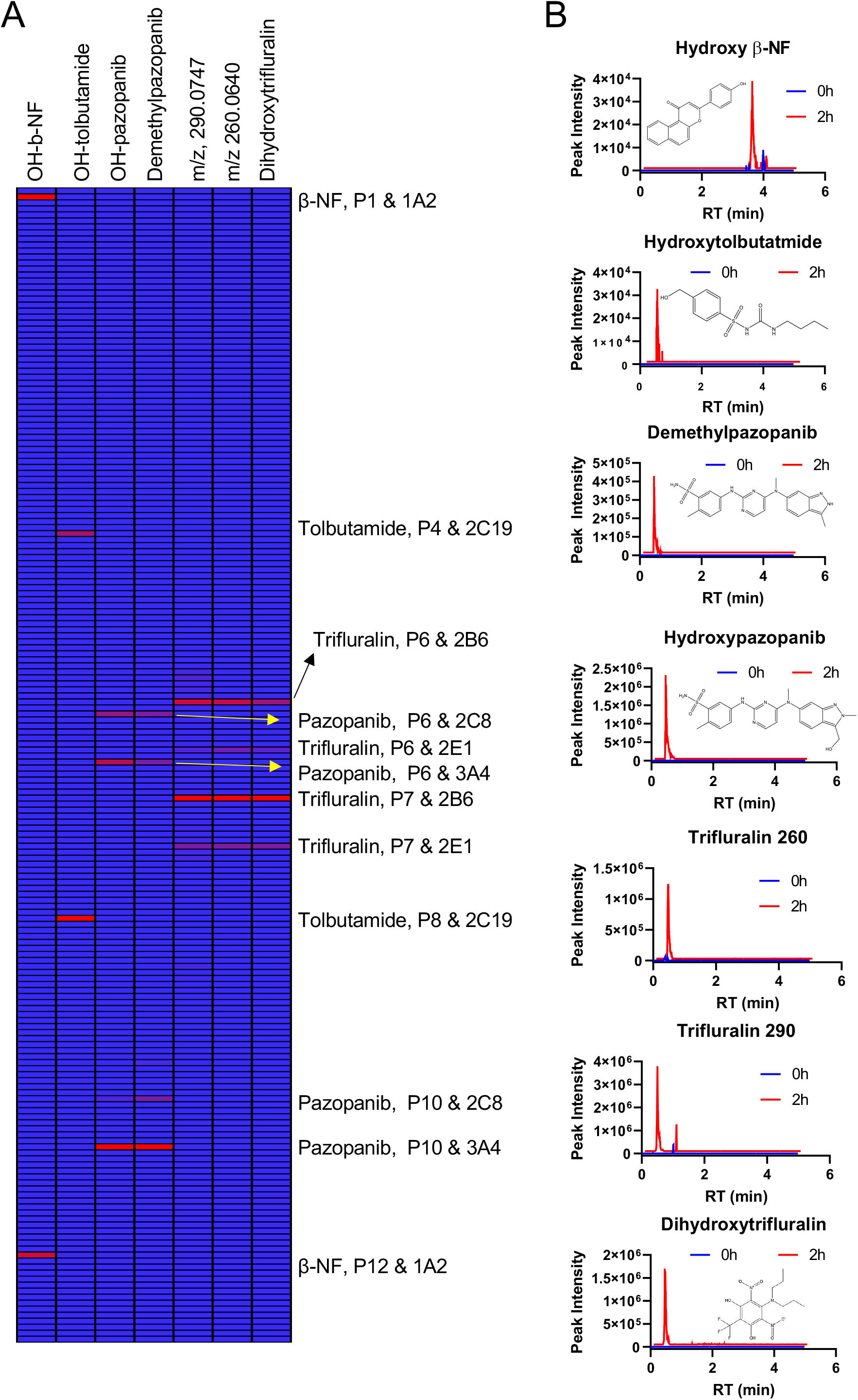
Results of the pooled chemical screen for 4 selected compounds. Each pool was incubated with P450-expressing cell lines for 0 and 2 h and quenched with ACN. Each sample was injected into HRMS as duplicate technical replicates and the signal was averaged. To increase the analytical stringency, a signal was removed if it was absent in one technical replicate. The same criteria as in Fig. 2 were used to select signals. A. Heat map of metabolites from 4 selected compounds. OH-tolbutamide was detected in C18 negative mode. The OH-tolbutamide signal was detected in pools 4 and 8 containing the parent tolbutamide after 2h incubation with the CYP2C19 cell line. Desmethyl and hydroxypazopanib signals were detected in pools 6 and 10 containing the parent pazopanib after 2h incubation with the CYP2C8 and CYP3A4 cell lines (HILIC positive mode). The OH-β-NF signal was detected in pools 1 and 12 containing the parent β-NF after 2 h incubation with the CYP1A2 cell line (C18 negative mode). Hydroxytrifluralin and two unidentified *m/z* were detected in HILIC positive mode. These signals were detected in pools 6 and 7 containing the parent trifluralin after 2 h incubation with the CYP2B6 and CYP2E1 cell lines. B. Chromatograms of metabolites. From top to bottom: OH-β-NF, OH-tolbutamide, desmethyl and hydroxypazopanib, unidentified trifluralin metabolite 260, unidentified trifluralin metabolite 290, dihydroxytrifluralin.

#### Examples of specific xenobiotics

Results for two well-studied drugs, tolbutamide and pazopanib, are shown in Fig. 6. Hydroxytolbutamide (Wester et al., 2000) was generated only by the CYP2C19 cell line in pools containing tolbutamide. Pazopanib is mainly metabolized by CYP3A4 and to a minor degree by CYP1A2 and 2C8 based on a previous *in vitro* study (Keisner and Shah, 2011). We detected hydroxypazopanib following incubations with CYP3A4 and CYP2C8 cells (and at significantly lower levels in one of the two CYP1A2 cell line incubations). Desmethyl pazopanib was generated only from CYP3A4 and CYP2C8 cell lines (Fig. 6). Similarly, hydroxy β-NF was detected only in incubations of the CYP1A2 cell line with pools containing β-NF (Pool 1 and Pool 12) shown here using C18 LC and negative ionization mode (Fig. 6). These data show that predicted metabolites were generated in a cell line specific manner from pooled mixtures, which verifies that a mixtures approach can be used with these cell lines to generate and detect xenobiotic metabolites.

#### Evidence for unidentified metabolites

Trifluralin is one of most widely used herbicides in the environment, whose health hazards are not known (https://www.epa.gov/sites/production/files/2016-09/documents/trifluralin.pdf). Identifying potential health risks associated with trifluralin exposure requires confident detection in human exposome studies. However, to the best of our knowledge, human P450 trifluralin metabolism is not characterized, which limits our ability to identify these exposures in human samples. Our experiments with trifluralin showed the generation of three features with *m/z* 260.0640, 290.0744, and 368.1059 by CYP2B6 and CYP2E1 cell lines (Fig. 6). The intensities at 290.0744, and 368.1059 were relatively high, while the *m/z* 260.0640 metabolite was generated from both cell lines at significantly lower levels. 368.1059 *m/z* is consistent with dihydroxytrifluralin, and MS/MS spectra are also consistent with this annotation (data not shown). However, the chemicals associated with 290.0744 *m/z* and 260.0640 *m/z* are currently unidentified.

#### Results from all 36 compounds

Having validated that the pooling strategy allowed for generation and detection of all predicted major metabolites from well-characterized compounds, we searched for potential metabolites of all 36 compounds based on several criteria: 1) the *m/z* feature should appear only in pools containing the associated parent compounds, 2) the same *m/z* feature should appear in both independent pools containing the same parent compound, 3) the *m/z* in both pools should show the same cell line specificity, and 4) the intensity for respective *m/z* features should show significantly lower levels or zero values in 0 h samples. Once we identified *m/z* based on these criteria, we annotated targeted metabolites using BioTransformer and a curated list of known mass differences caused by xenobiotic transformations (see Materials and Methods). Where possible, we confirmed reactions from literature searches.

Table 1 summarizes the cell-line dependent production of metabolites from the 36 compounds detected by HILIC positive ESI. Supplemental Table 1 shows the metabolite generation analyzed on C18 negative ESI. We identified many expected metabolites and hundreds of other *m/z* features which could represent uncharacterized metabolites generated from various compounds in a cell line specific manner. From 36 compounds, these cell lines generated a total of 825 unidentified mass spectral signals in HILIC positive ESI mode, suggesting that this high-throughput approach could be applied systematically to identify uncharacterized biotransformation products of drugs and environmental chemicals that are generated by specific human CYP enzymes.

## Discussion

Here, we demonstrate the generation of Huh7 hepatoma cell lines with stable expression of catalytically active, C-terminally-tagged P450 enzymes and demonstrate their substrate-specific P450 activities. By combining targeted mass spectral searching for known and expected metabolic products with powerful non-targeted LC-HRMS methods, we obtained evidence for a large number of uncharacterized metabolic products from common xenobiotics. We further showed that LC-HRMS has sufficient specificity and sensitivity to support a mixtures approach for metabolite detection. Specifically, incubation of six compounds together in each cell line increased analytical throughput by a factor of six, helping overcome throughput on the mass spectrometer as a bottleneck to large-scale characterization of xenobiotic metabolism. Using this approach, we show that these cell lines generate metabolites from the mixtures in a cell line and substrate-specific manner, allowing for specific P450 enzyme-substrate pairs to be identified.

We have previously shown that S9 enzymes can be used to provide a high-throughput approach to generate Phase I and II metabolites to identify xenobiotic exposures in humans (Liu et al., 2021). The P450 cell lines provide a complementary workflow for simultaneous generation of Phase I, CYP-dependent, metabolites of xenobiotics, and subsequent identification of active enzyme-substrate pairs. This capability to link substrates and specific metabolites with specific P450 cell lines could potentially be used for metabolic phenotyping studies in humans. This study shows that these P450 cell lines generate metabolites that are detected in human and experimental samples but not generated by the liver S9 supernatants. For example, bupropion hydroxylation by CYP2B6 is well documented (Hsyu et al., 1997; Hesse et al., 2000). Our data show that bupropion is hydroxylated in a CYP2B6 cell line dependent manner and that this specific cell line also generated dihydroxybupropion (RT 1.39 min), which was not detected previously using the S9 system (data not shown). Petsalo et al., (2007) first identified dihydroxybupropion in human urine samples using LC/MS/MS (Petsalo et al., 2007). Our data suggest that dihydroxybupropion is produced mainly through the activity of CYP2B6.

Other results show that the P450-expressing cells have activities expected for specific P450 enzymes. Omeprazole is metabolized mainly by CYP2C19 and CYP3A4 (Yamazaki et al., 1997; Li et al., 2005) and detected products as well as relative intensities of products are consistent with previous studies (Li et al., 2005; Ryu et al., 2014) showing selective hydroxylation of omeprazole by CYP2C19 and CYP3A4. Similarly, our results showing the deethylation of ethoxyresorufin by CYP1A2 cells, coumarin hydroxylation by CYP2A6 cells (Ghosal et al., 2003) and dextromethorphan demethylation by CYP2D6 cells (Yu and Haining, 2001) are consistent with known specificities of these enzymes.

These cell lines have several other possible utilities. The robust metabolite generation suggests that they could be used as an alternative to other approaches, e.g. baculosomes, for enzyme phenotyping and prediction of drug-drug interactions. In principle, the approach can be extended to develop additional P450-expressing lines and cells expressing UDP-glucuronosyltransferases and sulfotransferases. The cell lines provide several advantages including 1) ease of propagation and preservation under liquid nitrogen; 2) use without expensive and unstable cofactors such as NADPH, 3’-phosphoadenosine-5’-phosphosulfate and uridine diphosphate-glucuronic acid; and 3) metabolic product formation in cell environment representative of the native cellular environment *in vivo*.

The cell lines can also be used to generate specific metabolites in quantities for more conclusive identification by MS/MS and NMR spectroscopy. For example, we identified three major metabolites of trifluralin in CYP2B6 cell lines (Fig. 6). One metabolite was proposed to be dihydroxytrifluralin from MS/MS spectra; however, two other metabolites could not be identified by their MS/MS fragmentation patterns. Batch cultures of the CYP2B6 cell line with trifluralin suggest that sufficient quantities of products can be produced for purification and analysis by NMR to elucidate structures (manuscript in preparation). In addition, these cell lines can be used for studies on the toxicity of a chemical and its metabolites, analogous to those of Chen et al. (2021) who used a P450-expressing HepG2 cell line to study liver toxicity by amiodarone and dronedarone.

We learned that a mixtures approach (6 substrates per cell line per incubation) could be used to increase throughput for these large-scale screens, reducing the number of samples to be analyzed by a factor of 1/6 in this case. To analyze 36 individual compounds in duplicate with 8 different cell lines at 0 and 24h would require 8 × 2 × 2 × 36 = 1152 samples for LC/MS/MS analysis. By pooling (grouping 6 compounds as one pool) and adding each compound into two different pools, we reduced the number of samples for LC/MS/MS analysis to 192, significantly decreasing expense and increasing throughput. Requiring that a mass spectral signal be generated by a P450 cell line with two different pools containing a given xenobiotic increased stringency for qualifying a feature as a metabolite from that xenobiotic. Thus, the pooling strategy can be used as an initial screen to identify enzyme-compound specific activity pairs and to guide subsequent metabolite generation and characterization.

This mixtures approach could be vulnerable to confounding effects of potential substrate inhibition or competition among xenobiotics in a given pool. The likelihood of such interactions is mitigated to some extent by including each chemical in two different pools, and in this particular screen we did observe some instances where the amount of metabolite produced in two different pools containing the same compound were different. However, the appearance of metabolite in the two different pools still enabled the identification of the metabolites and the enzyme(s) that produced them. One could overcome this potential disadvantage using a “post-pooling strategy” in which individual chemicals are incubated with each cell line and then cell lysates and media are pooled for LC-HRMS analysis. However, this strategy requires more biological incubations, and samples may need to be further concentrated to generate similar levels of signal as the pre-pooling strategy described here. Additional limitations are that impure xenobiotic preparations can complicate interpretations with a mixtures approach, and reactive products from one xenobiotic could react with other xenobiotics. While the combination of xenobiotics in different pools protects against the latter complication, the problem of impurities is always present in drug metabolism research. On a positive side, the approach as developed is well suited for systematic P450 metabolism studies of drug formulations and compounded drug mixtures.

In conclusion, cell lines expressing xenobiotic enzymes can be used to metabolize xenobiotic mixtures to accelerate characterization and identification of xenobiotic metabolites for human high-resolution metabolomic studies. Our platform can be applicable for various additional research areas such as drug metabolism, drug-drug interactions, and mechanistic toxicology.

## Abbreviations

ACN: acetonitrile
β-NF: β-naphthoflavone
C18: reverse phase chromatography
CLZ: chlorzoxazone
DMEM: Dulbecco’s Modified Eagle Medium
ESI: electrospray ionization
G418: Geneticin
HILIC: hydrophilic interaction liquid chromatography
HRMS: high resolution mass spectrometry
LC: liquid chromatography
P450: cytochrome P450
PS: penicillin/streptomycin
RT: retention time

## Acknowledgements

Not applicable

## Author contributions

*Participated in research design:* Lee, Liu, Singer, Miller, Li, Jones, Morgan

*Conducted experiments:* Lee, Liu, Singer

*Performed data analysis:* Lee, Liu, Singer

*Wrote or contributed to the writing of the manuscript:* Lee, Liu, Singer, Miller, Li, Jones, Morgan

## Footnotes

This work was supported by the National Institutes of Health of Environmental Health Science [Grants U2C ES030163, 1P30ES019776].

This work has been deposited on the preprint server https://www.biorxiv.org.

## Notes

### Competing Interest Statement

The authors have declared no competing interest.

