## Supplemental Figs and Table for "High-throughput Production of Diverse Xenobiotic Metabolites with P450-transduced Huh7 Hepatoma Cell Lines"

A

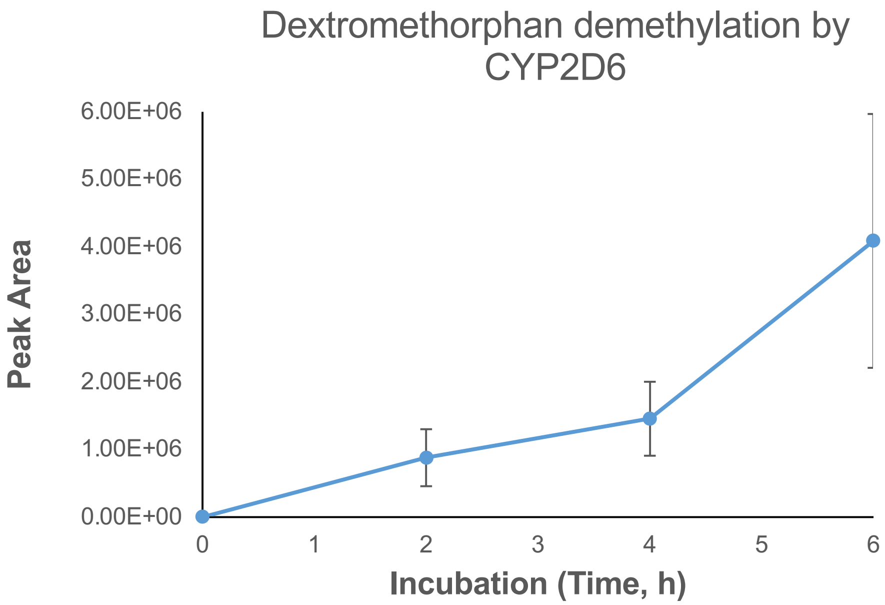


B

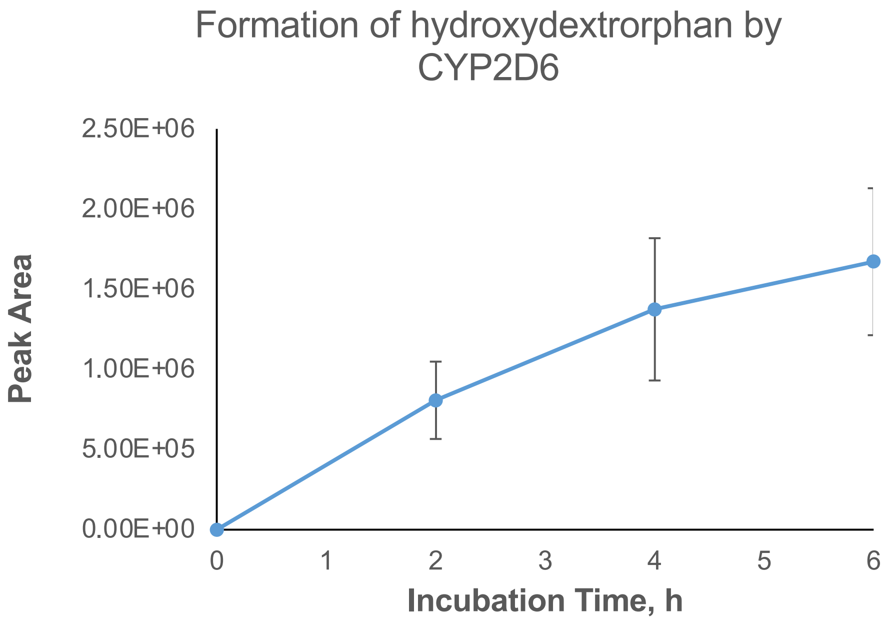


**Supplemental Fig. 1**. Cell line and substrate specificity. To examine substrate specificity of these cell lines, all cell lines were incubated for the indicated times with a mixture of 7 substrates, and then the whole contents (media and cells) of each well were analyzed by LC-HRMS on the Orbitrap ID-X Tribrid MS. A. Demethylation of dextromethorphan by 2D6 B. Formation of hydroxydextrorphan by CYP2D6 following demethylation of dextromethorphan. Values are means ± SD of three individual wells from the same cell preparation.

Pool 36 chemicals, each in two unique pools

| Pool number | P1 | P2 | P3 | P4 | P5 | P6 | P7 | P8 | P9 | P10 | P11 | P12 |
|---|---|---|---|---|---|---|---|---|---|---|---|---|
| Chemical numbers | C1 | C7 | C13 | C19 | C25 | C31 | C1 | C2 | C3 | C4 | C5 | C6 |
| | C2 | C8 | C14 | C20 | C26 | C32 | C7 | C8 | C9 | C10 | C11 | C12 |
| | C3 | C9 | C15 | C21 | C27 | C33 | C13 | C14 | C15 | C16 | C17 | C18 |
| | C4 | C10 | C16 | C22 | C28 | C34 | C19 | C20 | C21 | C22 | C23 | C24 |
| | C5 | C11 | C17 | C23 | C29 | C35 | C25 | C26 | C27 | C28 | C29 | C30 |
| | C6 | C12 | C18 | C24 | C30 | C36 | C31 | C32 | C33 | C34 | C35 | C36 |

Incubate each pool with
8 CYP cell lines for 0 and 24 h

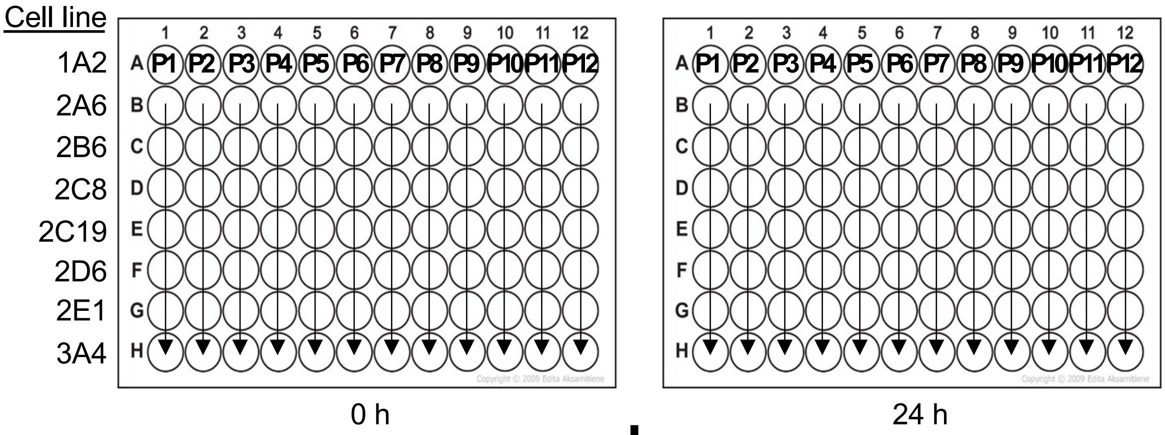


Quench reaction with 3 volumes acetonitrile
Extract sample for LC/MS/MS analysis

**Key to chemicals**
C1, acenapthene; C2, fluoranthene; C3, 1,1,2,2-trichloroethane; C4, azinphosmethyl; C5, tribufos; C6, β-naphthoflavone; C7, chlorzoxazone; C8, 1,2-diphenylhydrazine; C9, N-nitrosodipropylamine; C10, dexamethasone; C11, bromodichloromethane; C12, 1,2-dichlorobenzene, C13, acetaminophen; C14, ethylbenzene,; C15, benzo[a]pyrene; C16, bupropion; C17, ethion; C18, vendetanivb; C19, naphthalene; C20, tolbutamide; C21, 2-methynaphthalene; C22, G418; C23, benzo[b]fluoranthene; C24, o,p'-DDD; C25, coumarin; C26, 2,4,6-trichlorophenol; C27, endrin aldehyde; C28, caffeine; C29, bis(2-hexylethyl)phthalate; C30, nicotine; C31, trifluralin; C32, estrone; C33, auranofin; C34, pazopanib; C35, crizotinib; C36, warfarin

**Supplemental Fig. 2**. Pooling strategy. A total of 36 compounds were grouped into 12 pools, with each pool containing a unique combination of 6 compounds and each compound being present in two separate pools.
B. Experimental design. After each pool incubated was incubated with 8 CYP-expressing cell lines for 0 and 2h on two 96-well plates, respectively, reactions were quenched with 3 volumes of ACN and then analyzed by LC-HRMS (Orbitrap Fusion™ Tribrid™ Mass Spectrometer)

Supplemental Table 1. Xenobiotic biotransformation products from CYP cell lines. Metabolites detected in C18-negative mode are shown. Targeted metabolites are named. Numbers in parentheses represent unidentified biotransformation products which were strongly associated with parent compounds and identified by the criteria stated in the text. (Pearson co-efficient >0.85).

| **CYP cell line** | **1A2** | **2A6** | **2B6** | **2C8** | **2C19** | **2E1** | **2D6** | **3A4** |
|---|---|---|---|---|---|---|---|---|
| Acenaphthene | | | | | | | | |
| Fluoranthene | 23 | 23 | 15 | 63 | 1 | 34 | | 22 |
| 1,1,2,2-Tetrachloroethane | 1 | | | | | | | |
| Azinphos methyl | | | | | 1 | | 1 | |
| s,s,s,-tributyl_phosphorotrithioate | | | | | 7 | | | 7 |
| b-naphthoflavone | 1 | | | | | | | 1 |
| Chlorzoxazone | 1 | 1 | 1 | 1 | 1 | hydroxy; 2 | 1 | 4 |
| 1,2-diphenyl hydrazine | | | | | | | | |
| N-Nitrosodipropylamine | | | | | | | | |
| Dexamethasone | 2 | 2 | 1 | 2 | 2 | 1 | 4 | |
| Bromodichloromethane | | | | | | | | |
| 1,2-dichlorobenzene | 1 | | | 1 | | | | |
| APAP | 1 | | | | | | | 1 |
| Ethylbenzene | 2 | | | | | | | |
| Benzo[a]pyrene | | 1 | | | | | | |
| Bupropion | 1 | | | | | | | |
| Ethion | | | | 1 | 7 | 1 | | 2 |
| Vandetanib | 3 | 3 | | 1 | | 2 | 1 | 1 |
| Naphthalene | | | | | | | | 2 |
| Tolbutamide | | | | | hydroxy; 1 | 1 | 3 | 2 |
| 2-Methylnaphthalene | | hydroxy | | | | | | |
| G418 | | | | | 2 | 1 | | |
| Benzo-b-fluoranthene | | | 1 | | | | 1 | |
| o,p,-DDD | | hydroxy; 1 | hydroxy, dihydrodiol; 22 | | | 12 | | |
| Coumarin | | hydroxy; 6 | | | | | | |
| 2,4,6-Trichlorophenol | | | | | | | | |
| Endrin aldehyde | 8 | 27 | 20 | 31 | 33 | 22 | 42 | 40 |
| Caffeine | 1 | | | | | | | |
| Bis-2-ethylhexyl-phthalate | | 1 | 1 | | | | 1 | |
| Nicotine | | | | | | | | |
| Trifluralin | | 1 | loss of propane and ketone formation; 76 | 3 | | loss of propane and ketone formation; 51 | | 6 |
| Estrone | | | | | | | | demethylation, hydroxy; 6 |
| Auranofin | de-acetylation; 1 | de-acetylation; 2 | | de-acetylation; 6 | de-acetylation; 5 | de-acetylation; 2 | de-acetylation; 9 | de-acetylation; 7 |
| Pazopanib | | | | hydroxy; 1 | 1 | | | demethyl; 1 |
| Crizotinib | | | | | 1 | 1 | 1 | |
| Warfarin | 3 | 2 | | 2 | hydroxy; 3 | | 4 | 1 |
